# Chromosome-scale *Daphnia magna* genome assembly resolves scaffolding discrepancies

**DOI:** 10.64898/2026.09.03.749101

**Authors:** Pascal Angst, Peter D. Fields, Christoph R. Haag, Dieter Ebert

## Abstract

*Daphnia magna* is a widely used model organism in ecological genomics, ecotoxicology, and evolutionary biology. While a few genome assemblies of this species of freshwater crustacean are available, disagreements in their chromosomal arrangements point toward scaffolding problems. We present the most contiguous chromosome-level genome assembly of *D. magna*, the first based on PacBio HiFi sequencing technology. The assembly spans 184.38 Mb and has a scaffold N50 of 12.4 Mb. Eight of ten chromosomal scaffolds consist of mainly two contigs, representing one chromosome arm each. Eleven of 20 telomeric regions are fully resolved. Using a genomic map and comparative genomics, we were able to not only resolve the problems of previous assemblies, but also to demonstrate that some of the scaffolding errors followed a systematic pattern, most likely resulting from mis-joins during Hi-C scaffolding. Additionally, we provide expression profiles for annotated genes derived from different *D. magna* RNA-seq datasets, enabling rapid assessment of predicted gene expression and identification of potential pseudogenes. The new genomic and transcriptomic resource enables comparative genomic studies on genome architecture, structural variation, and gene family evolution in this important model organism.

**Significance Statement:** Genome assemblies of different individuals of the same species vary as a result of natural genetic variation and assembly errors but distinguishing between these two sources of variation can be challenging. We generated an updated *D. magna* genome using long, highly-fidelity DNA sequences and a genetic map, revealing that previous assemblies contain systematic errors, including inverted chromosome arms likely due to mis-joins from the widely used Hi-C scaffolding method. The new genome enables identifying real genetic differences among the many already sequenced *D. magna* individuals from across the species range, allowing to investigate how natural selection shapes large-scale genetic variation, for example, involved in resistance to stressors and parasites.

## Introduction

*Daphnia magna* is a model system for ecological and evolutionary genomics (Ebert, 2022). The freshwater crustacean plays an important role in aquatic ecosystems as a central link in the food web, particularly through its filter feeding activity (Lampert, 2025). As an experimental model, it offers short generation times, clonal reproduction enabling genetic replication, and well-established protocols to test responses to environmental stressors (OECD, 2004, 2012). Genomic studies of *D. magna* and its parasites have advanced our theoretical and empirical understanding of host–parasite interactions and coevolution by combining natural as well as experimental parasite infections with whole-genome sequencing (Bento et al., 2017; Dexter et al., 2023).

Previous studies of the *D. magna* genome revealed that parasite resistance genes occur in regions with extensive structural variation termed large structural polymorphisms (LSPs) (Dexter et al., 2026; Naser-Khdour et al., 2024). Recently, the Pasteuria Resistance Complex (PRC) on chromosome 4 and a multi-megabase LSP on chromosome 5 were shown to have immunity-related signatures of long-term balancing selection from local to species distribution-wide geographic scales and to undergo rapid frequency shifts during parasite epidemics (Bourgeois et al., 2021; Cornetti et al., 2024; Dexter et al., 2026). Characterizing such structural variants and investigating their evolution requires chromosome-scale genome assemblies. While a few genome assemblies of *D. magna* were scaffolded to chromosomes (Byeon et al., 2022; Chaturvedi et al., 2023; Lee et al., 2019), they vary in contiguity and chromosomal arrangement.

Here we present a chromosome-scale genome assembly for *D. magna* that addresses these inconsistencies by combining new high-fidelity long-read sequencing and existing high-density linkage maps (Dukić et al., 2016; Routtu et al., 2010, 2014). We also provide transcript quantification derived from RNA-seq data by combining datasets from diverse projects. This allows us to expand genomic and transcriptomic resources for reliably detecting and analyzing structural variation and gene expression in *D. magna*.

## Materials and Methods

### Sample and sequencing

The sequenced *D. magna* genotype originated from an individual female collected from a small rock pool on a skerry island near the Tvärminne Zoological Station in South-Western Finland (latitude: 59°49’58’’□N, longitude: 23°15’45’’ E). This female was propagated in the laboratory via parthenogenetic (clonal) reproduction and then sexually selfed by within-clone mating for three generations and kept as an iso-female line (clone ID: FI-Xinb3). High molecular weight DNA extraction using Genomic-tips (QIAGEN, Hilden, Germany) as described in Naser-Khdour et al. (2024). PacBio (Menlo Park, CA, USA) low DNA input SMRTbell library preparation and HiFi sequencing on a Sequel II machine at the Lausanne Genomics Technologies Facility (GTF, UNIL, Switzerland) generated 20.5 Gb of sequencing data (107× coverage, read N50: 10.9 Kb) of the FI-Xinb3 clone.

### Assembly

PacBio HiFi reads were assembled using hifiasm v.0.25.0 (Cheng et al., 2021). Non-target contigs were filtered from the assembly using the NCBI Foreign Contamination Screen v.0.5.5 and BlobTools v.1.1.1 (Laetsch & Blaxter, 2017), which relied on results from BLAST+ v.2.17.0 BLASTn (Camacho et al., 2009) searches against NCBI nt and Diamond v2.1.23 blastx (Buchfink et al., 2021) searches with parameters -F 15 -b4 -c1 against UniProt Reference Proteomes. We removed haplotypic duplication using purge_dups v.1.2.6 (Guan et al., 2020) with previously generated long-read sequences of the same clone (NCBI database, BioProject ID: PRJNA624896, Cornetti et al. (2024)). We scaffolded our assembly using the *D. magna* high-density genetic map (Dukić et al., 2016), specifically by aligning it to the ordered and oriented v.2.4 FI-Xinb3 assembly (NCBI database, Assembly name: daphmag2.4, GenBank assembly accession: GCA_001632505.1, BioProject ID: PRJNA298946, Daphnia Genomics Consortium) using minimap2 v.2.30 (Li, 2021) with the -ax asm5 parameter. Contig joins were filled with 100 Ns, unless filled with read sequence using TGS-GapCloser v.1.2.1 (Xu et al., 2020). Assembly completeness was assessed using BUSCO v.6.1.0 (Tegenfeldt et al., 2025) against the arthropoda_odb12.2 (Creation date: 2026-05-13) lineage and visualized using BlobToolKit2 v.4.4.5 (Challis et al., 2020).

### Annotation

Gene annotations were transferred from the v.3.0 assembly of the FI-Xinb3 genome (Zenodo DOI: https://doi.org/10.5281/zenodo.11270426, Cornetti et al. (2024)) based on sequence similarity (Table S1) using Liftoff v.1.6.3 (Shumate & Salzberg, 2021) with -chroms, -copies, and -polish flags. Repetitive elements were identified using RepeatModeler2 v.2.0.7 with the LTR pipeline (Flynn et al., 2020) to build a species-specific repeat library, followed by RepeatMasker v.4.2.3 (Smit et al., 2013). Telomeric repeats were identified by (TTAGG)n motifs at scaffold termini. Annotations were visualized with the packages RIdeogram v.0.2.2 (Hao et al., 2020), Biostrings v.2.76.0 (Pagès et al., 2025), and GenomicRanges v.1.60.0 (Lawrence et al., 2013) in R v.4.5.3 (R Core Team, 2026)

### *Daphnia* genome assemblies for comparison and synteny analysis

Available chromosome-level *D. magna* genome assemblies were downloaded from NCBI for comparative analyses: LRV0_1 (NCBI database, Assembly name: UOB_LRV0_1, GenBank assembly accession: GCA_030254905.1, BioProject ID: PRJNA777104, Chaturvedi et al. (2023)), NIES (ASM2063170v1.1, GCA_020631705.1, PRJNA738190, Byeon et al. (2022)), SK or KIT (ASM399081v1, GCA_003990815.1, PRJNA490418, Lee et al. (2019)). The Daphnia pulex NCBI reference genome assembly KAP4 (ASM2113471v1, GCA_021134715.1, PRJNA777597, Ye et al. (2025)) was also downloaded for synteny analysis using the GENESPACE v.1.3.1 (Lovell et al., 2022) R package.

### Expression profiling

RNA-seq datasets from the STRESSFLEA consortium and transcriptomic studies of *D. magna* were used for transcript quantification (Molinier et al., 2019; Orsini et al., 2016). The STRESSFLEA dataset consists of 67 libraries from three *D. magna* genotypes (FI-Xinb3, DE-Iinb1, and an F2 recombinant cross of these genotypes) exposed to twelve environmental stressors (NCBI database, BioProject ID: PRJNA284518). RNA-seq reads were quality-trimmed using fastp v.1.0.1 (Chen, 2023) and quantified against the decoy-aware reference transcriptome using Salmon v.1.10.3 (Patro et al., 2017) with -- keepDuplicates and --validateMappings flags. The reference transcriptome consisted of transcripts extracted from the genome annotations using GffRead v.0.12.7 (Pertea & Pertea, 2020) and filtered to retain sequences of at least 200 bp using SeqKit2 v.2.10.1 (Shen et al., 2024). The Salmon software reports expression as transcripts per million (TPM). Genes with TPM greater than one in at least two RNA-seq libraries were classified as expressed.

## Results

### Assembly statistics

Our chromosome-scale genome assembly of *D. magna*, the first based on PacBio HiFi sequencing technology, spanned 184.38 Mb, with a scaffold N50 of 12.4 Mb and contig N50 of 6.4 Mb. Its contiguity and completeness surpassed those of all previous *D. magna* assemblies. The contig N50 was at least four times greater than that of any prior assembly, with more sequence content anchored into the ten chromosomal scaffolds (Table 1). The chromosomal scaffolds comprised 31 contigs: Eighteen of these represented chromosome arms, while the remaining 13 shorter contigs represented either half-arms or regions of the centromere or telomere. Eleven chromosome ends had (TTAGG)n telomere repeats at or near the terminus, compared to none in all previous assemblies. Three chromosomes were assembled telomere-to-telomere, except for partially missing centromere sequence. These are the first telomere-to-telomere scaffolds of *Daphnia*. The assembly had an area under the Nx curve (auN) of 11.2 Mb and a snail score of 0.61 (scores > 0.6 represent high-quality assemblies; Figure 1, Challis & Blaxter (2026)). BUSCO analysis indicated 98.5% complete BUSCOs (97.1% single-copy, 1.4% duplicated). The genome contained 50.79% (93.65 Mb) repetitive DNA (Table 2), with repeat enrichment in (peri)centromeric and (sub)telomeric regions (Figure 2). The most abundant repeat class at 32.93% of the genome was LTR retrotransposons, mostly from the Ty3-gypsy family, while 6.45% of all repeats likely represented *Daphnia*-specific or other repeat families unknown to the Dfam and RepBase databases (Table 2).

**Table 1:** Statistics of the focal and previous *D. magna* genome assemblies.

| Clone ID | FI-Xinb3 | LRV0_1 | NIES | SK / KIT |
| --- | --- | --- | --- | --- |
| Assembly size (Mb) | 184.38 | 130.00 | 161.47 | 122.95 |
| Chromosomal scaffolds | 10 | 10 | 10 | 10 |
| Length of scaffolds (Mb) | 141.01 | 122.62 | 133.28 | 99.50 |
| Scaffold N50 (Mb) | 12.40 | 11.90 | 12.55 | 10.13 |
| Contig N50 (Mb) | 6.41 | 1.29 | 1.51 | 0.01 |
| complete BUSCOs (%) | 98.5 | 98.0 | 99.2 | 98.5 |
| single copy BUSCOs (%) | 97.1 | 97.2 | 93.9 | 97.7 |
| Telomeres resolved | 11/20 | 0/20 | 0/20 | 0/20 |
| Publication | new | Chaturvedi et al. (2023) | Byeon et al. (2022) | Lee et al. (2019) |

**Table 2:**
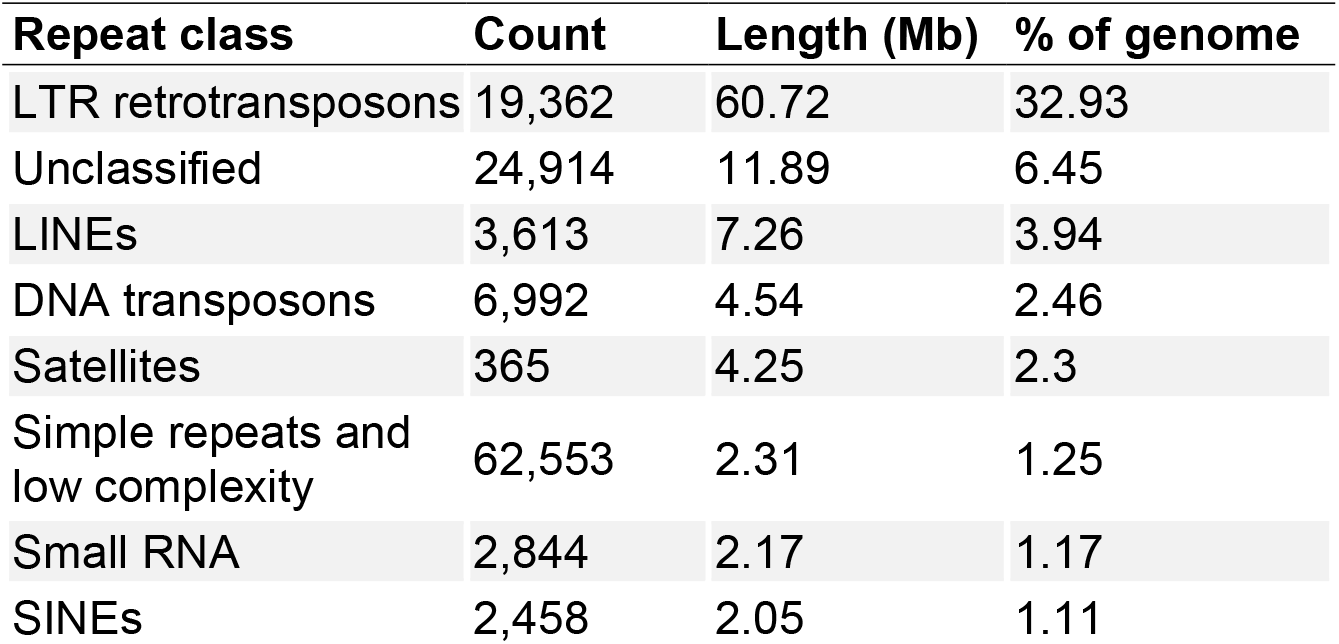
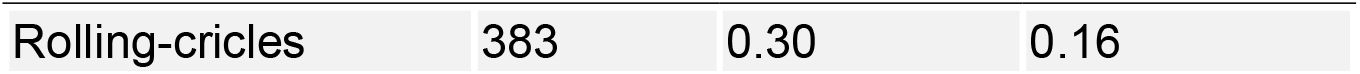
Summary of repeat annotation.

**Figure 1:**
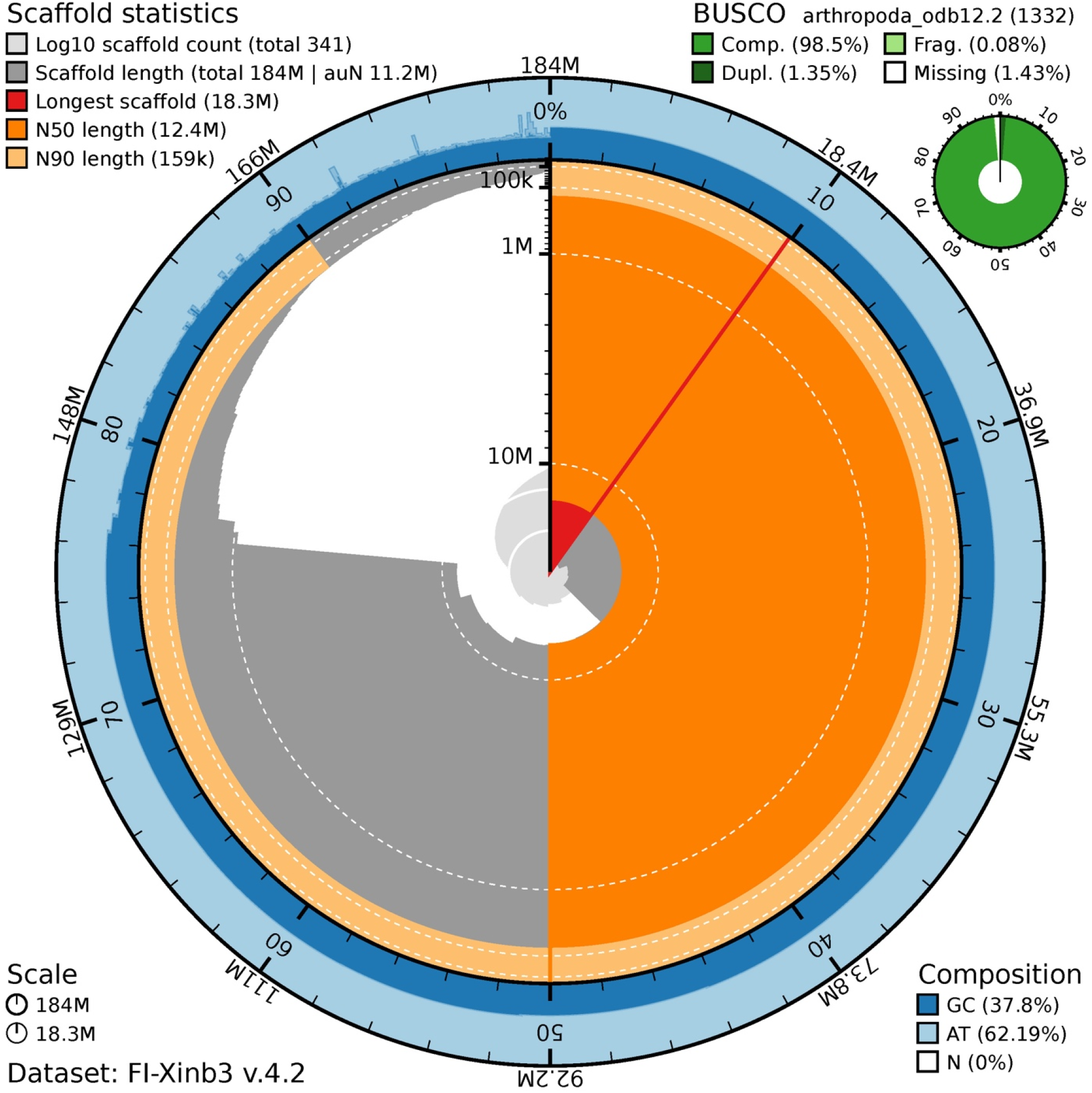
Snail plot of the *D. magna* FI-Xinb3 genome assembly, indicating high quality and providing a genome-size-independent summary of assembly metrics. The spiral (center) represents the cumulative length in Mb colored by longest scaffold, scaffold N50, and scaffold N90. The outer ring (blue) shows the GC content distribution (%). The small circle shows BUSCO gene completeness (%). Note: The sequences much shorter than the ten near-chromosomal scaffolds are unplaced contigs.

**Figure 2:**
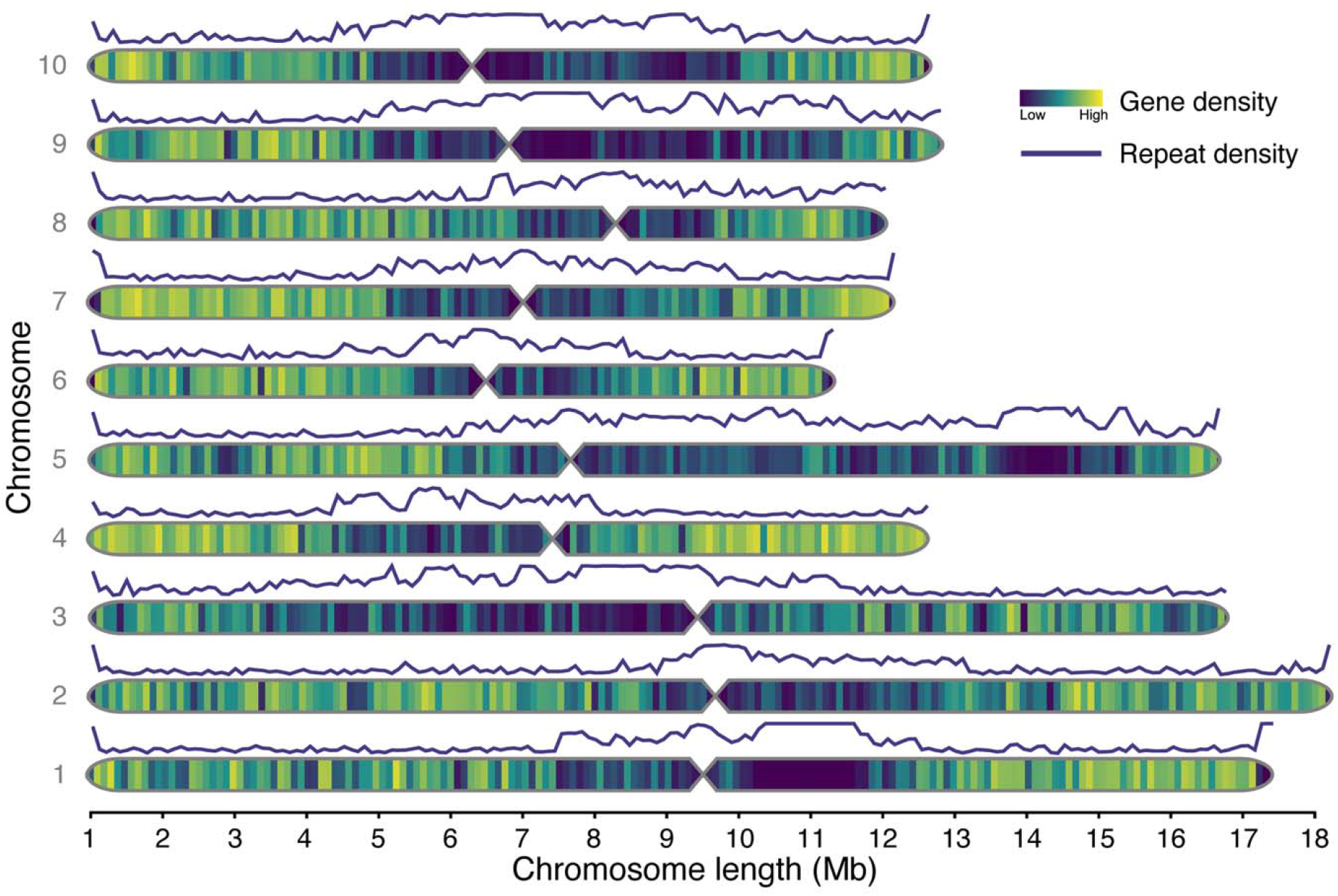
Ideogram of the *D. magna* FI-Xinb3 genome. The ten chromosomes are overlaid with heatmaps showing the density of expressed genes as a fraction of genic bases per 100 kb window, revealing gene depletion in repetitive regions. Lines above each chromosome indicate repeat density (fraction of repetitive bases per 100 kb window), showing repeat enrichment in (peri)centromeric and (sub)telomeric regions. Hourglasses spanning 400 Kb between chromosome arms mark approximate centromere locations inferred from contig joins. Chromosome numbers were adopted from Dukić et al. (2016). Note: In previous ideograms of the *D. magna* FI-Xinb3 genome (based on assembly v.3.1), chromosome arms, including 1R, 2R, 2L, 3L, 5R, 7R, 8R, and 9R, were shown in reverse orientation (i.e., Dexter et al., 2026; Otte et al., 2024; Naser-Khdour et al., in review).

### Synteny analysis

Our new genome revealed several scaffolding errors in previously published *D. magna* genomes when we compared all assemblies in a synteny analysis based on large-scale collinearity of orthogroups (Figure 3). The Hi-C-scaffolded LRV0_1 assembly exhibited incorrect intra-chromosomal joins, including multiple chromosome arm inversions that positioned (peri)centromeric sequences near scaffold ends rather than in their center, and vice versa. In contrast, the FI-Xinb3, NIES, and SK assemblies showed expected chromosomal collinearity, though there were some rearrangements that may be attributed to short contigs underlying the NIES and SK assemblies, complicating their scaffolding.

**Figure 3:**
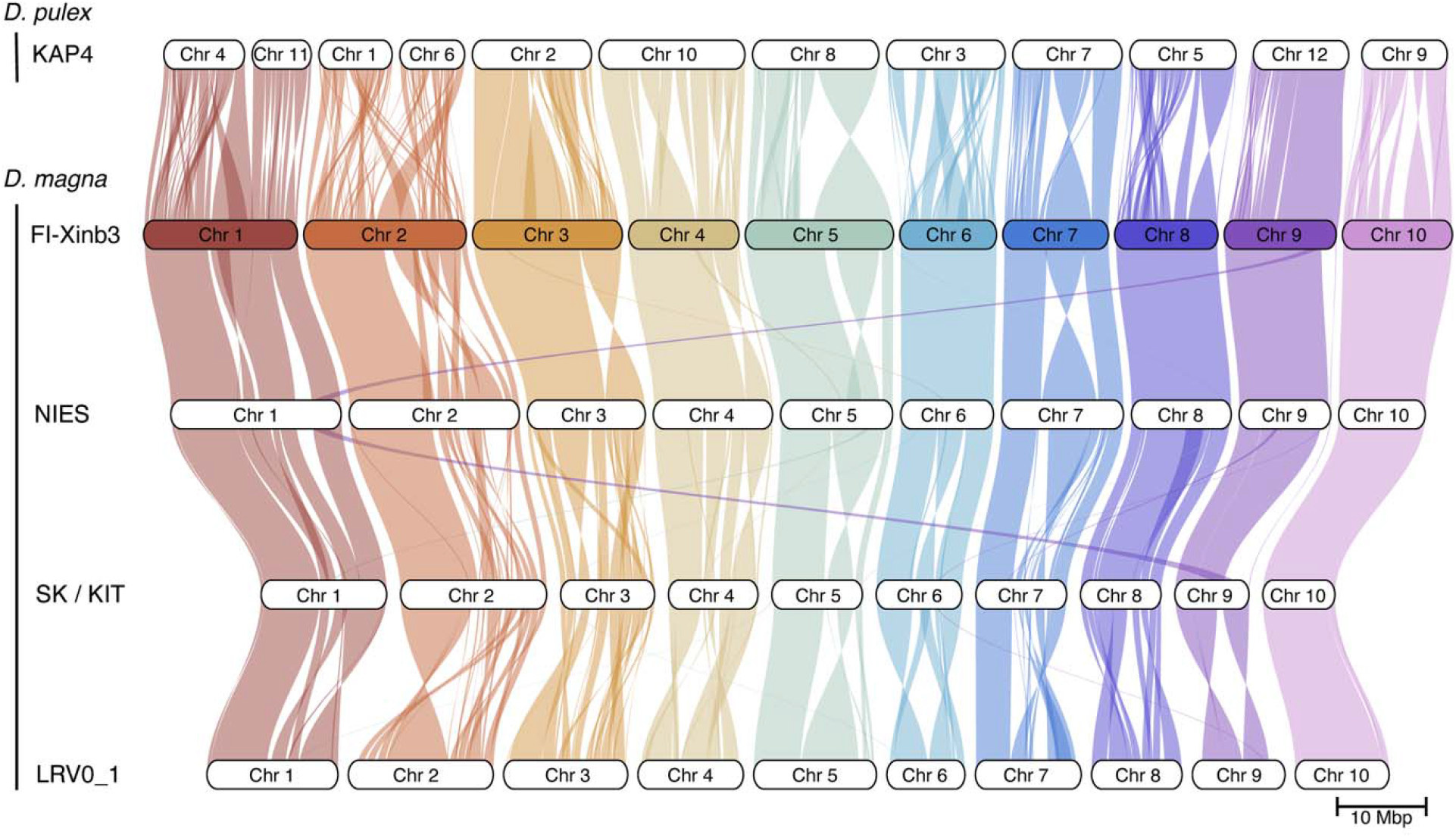
Riparian plot showing collinearity and rearrangements among *Daphnia* genome assemblies. Chromosomal scaffolds of *D. magna* assemblies (FI-Xinb3, NIES, SK, LRV0_1) and the *D. pulex* (KAP4) outgroup are represented as horizontal bars connected by ribbons, colored by *D. magna* chromosome assignment. Twisted ribbons to the LRV0_1 Hi-C-scaffolded assembly indicate chromosome arm inversions relative to the other assemblies.

### Genome annotation and transcript quantification

Of 31,070 predicted protein-coding genes, transferred from the previous v.3.0 annotation of the FI-Xinb3 clone, 21,148 genes (68.07%) showed expression greater than one TPM in at least two RNA-seq libraries of *D. magna* (Table S2). Contrarily, 316 genes showed zero expression across all libraries, representing candidates for pseudogenization. Genes with expression below our stringent threshold might show expression due to a few miss-mapped transcripts and are unreliable for identification of expressed genes versus pseudogene candidates. These potential false positives tended to occur in repetitive regions of the genome (compare Figures 2 and S1).

## Discussion

We present a highly improved assembly of the *D. magna* genome with 18 of 20 chromosome arms assembled into single contigs and eleven fully resolved telomeres, representing definitive chromosome boundaries. Comparison with previous assemblies revealed large-scale sequence differences primarily caused by mis-scaffolding: Short contigs comprised accurate scaffolding in the NIES and SK assemblies, while the more recent LRV0_1 assembly contained systematic Hi-C scaffolding mis-joins, particularly chromosome arm inversions that misplaced chromosome ends in chromosomal interiors, and vice versa.

Hi-C mis-joins, particularly inversions, are common because contig orientation is inferred by maximizing Hi-C contact support between contig ends (Dudchenko et al., 2018; Ghurye et al., 2019). This support is weak or ambiguous for short contigs and in repetitive regions, such as those close to centromeres. Recent sequencing technologies that produce longer primary contigs and improved Hi-C scaffolding methods that give more weight to contig end interactions have mitigated this problem (Lin et al., 2024; Wang et al., 2022; Zhou et al., 2023). Nevertheless, manual curation through contact map inspection and ideally additional orthogonal validation (i.e., optical maps, linkage maps, ultra-long reads) remain crucial for identifying and rectifying Hi-C mis-joins (Kadota et al., 2020; Marcolungo et al., 2023). Sequence characteristics of (peri)centromeric or (sub)telomeric regions, including repeat composition and GC content, can be used to orient contigs and scaffolds that represent chromosome arms when validation data is limited. Through high-fidelity long-read sequencing, validation against the genetic maps, and identification of telomere repeats, we provide a reliable reference with strongly reduced scaffolding errors, enabling accurate structural variant detection.

Our gene expression quantification across diverse RNA-seq libraries from different conditions and our repeat annotation provide additional resources for studying the evolution of gene families and transposable elements, particularly valuable for the fast-evolving, repeat-and immune gene family-rich LSP regions of the *D. magna* genome. We found evidence for the expression of 31,070 predicted protein-coding genes, including 21,148 with robust evidence. This is consistent with earlier suggestions that the *Daphnia* genome contains more than 30,000 genes (Colbourne et al., 2011).

## Conclusion

Modern genome assembly methods, including Hi-C scaffolding, substantially improve assembly contiguity but can introduce systematic errors even with careful curation, a problem documented across diverse taxa (Dudchenko et al., 2018; Ghurye et al., 2019). We demonstrate a generalizable approach integrating high-fidelity long-read sequencing with validation through genetic maps or comparative genomics to identify and correct such artifacts. This approach enables to distinguish real structural variation from technical artifacts, thereby facilitating studying how structural variation is shaped by adaptive and stochastic evolutionary processes using pangenomic, comparative genomic, and population genomic approaches in *D. magna* and other systems with extensive resequencing data.

## Data Availability

Raw data and the assembled genome will be deposited in the NCBI database (BioProject ID: PRJNA624896) and the predicted protein sequences plus expression profiles are available at Zenodo (Zenodo Preview: https://zenodo.org/records/20810573?preview=1&token=eyJhbGciOiJIUzUxMiJ9.eyJpZCI6IjUxN2I5YTk1LTQ5Y2EtNDk4OC1iYjk2LWU1NDZkNGVlYjYzMiIsImRhdGEiOnt9LCJyYW5kb20iOiIwMmQ1MmVhN2ZhYjg2MjFhYjMxNDc3NzQ4NWNjY2U5MiJ9.1GSfwrOp1ygMb_V2Nl4tSuL8DdnVoViUU9hS6lAwFKNYOefRJznMH-UAB1eSiXeSH6q_Lh1DC_3cYh_BPDJPNA).

## Acknowledgments

Claude Sonnet 4.5 (Anthropic) was used to provide proofreading and grammar suggestions.

---

**Figure S1:**
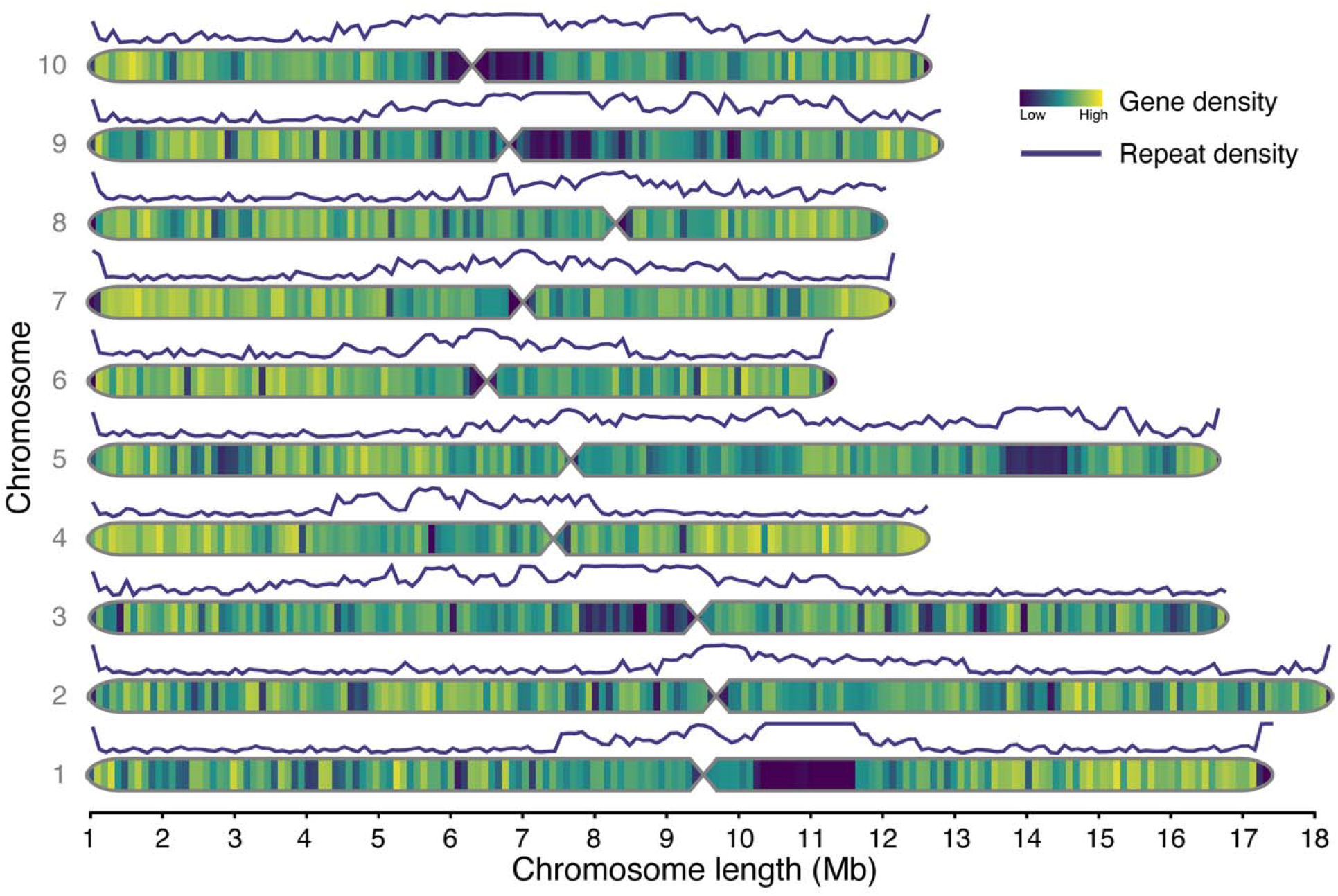
Ideogram of the *D. magna* FI-Xinb3 genome with all predicted genes of the transferred v.3.0 annotation. The ten chromosomes are overlaid with heatmaps showing the density of predicted genes as a fraction of genic bases per 100 kb window. Lines above each chromosome indicate repeat density (fraction of repetitive bases per 100 kb window). Hourglasses spanning 400 Kb between chromosome arms mark approximate centromere locations inferred from contig joins.

**Table S1: Matching assembly contigs of the focal and the v.3.0 FI-Xinb3 assembly based on sequence identity and coverage**.

**Table S2: Transcript quantification across *D. magna* RNA-seq datasets**. Expressed_any indicates TPM greater than 0 in at least one RNA-seq library and expressed_strict indicates TPM greater than 1 in at least two RNA-seq libraries.

